# Eosinophils are an endogenous source of IL-4 during filarial infections and contribute to the development of an optimal T helper 2 response

**DOI:** 10.1101/2023.10.26.564180

**Authors:** Cécile Guth, Pia Philippa Schumacher, Archena Vijayakumar, Hannah Borgmann, Helene Balles, Marianne Koschel, Frederic Risch, Benjamin Lenz, Achim Hoerauf, Marc P. Hübner, Jesuthas Ajendra

## Abstract

Interleukin-4 (IL-4) is a central regulator of type 2 immunity, crucial for the defense against multicellular parasites like helminths. This study focuses on its roles and cellular sources during *Litomosoides sigmodontis* infection, a model for human filarial infections. Our research uncovers eosinophils as a major source of IL-4, especially during the early phase of filarial infection. Using dblGATA mice lacking eosinophil and subsequently eosinophil-derived IL-4, we reveal their profound impact on the Th2 response. Lack of eosinophils impact Th2 polarization and resulted in impaired type 2 cytokine production. Surprisingly, eosinophil deficiency had no impact on macrophage polarization and proliferation as well as on antibody production. These findings shed new light on IL-4 dynamics and eosinophil effector functions in filarial infections.

**AUTHOR SUMMARY:** Filarial nematodes can cause severe diseases like onchocerciasis and lymphatic filariasis, posing a significant public health challenge in tropical regions, putting over a billion people at risk. The WHO categorizes these infections as neglected tropical diseases and aims to eliminate onchocerciasis transmission and lymphatic filariasis as a public health issue by 2030. To achieve this goal, we need a better understanding of the protective immune responses involved. Eosinophils have been identified as a key immune cell type in the well-established murine model for filarial infection, *Litomosoides sigmodontis*. However, their precise roles and interactions with other components of the type 2 immune response remain unclear. Our study reveals that eosinophils play a crucial role as a primary source of interleukin-4, the central cytokine in type 2 immunity. By using dblGATA mice, we found that the absence of eosinophils resulted in a reduced T helper 2 response but did not impact the alternative activation of macrophages or antibody production. In summary, our research uncovers an underappreciated function of eosinophils and their significant influence on type 2 immune responses.

## INTRODUCTION

Interleukin-4 (IL-4) plays a pivotal role in orchestrating the immune response associated with type 2 immunity, primarily aimed at defending against large multicellular parasites like helminths^1^. IL-4 exerts its influence across a spectrum of innate and adaptive immune cells. One of its critical functions is to guide the differentiation of T cells into T helper 2 (Th2) cells^2^. IL-4 activates the transcription factor STAT6 leading to the expression and activation of GATA3, the master regulator of Th2 cells. These Th2 cells then produce not only IL-4 but also the other signature cytokines IL-5 and IL-13^2^. Together with IL-13, IL-4 engages the IL-4 receptor, triggering an alternate activation pathway in macrophages^3^. This activation is a cornerstone of wound healing and tissue repair during type 2 immune responses^4^. IL-4 also plays a crucial role in B cell activities, including class switching and the production of immunoglobulin E (IgE)^5^. Furthermore, it dampens inflammatory responses, acting as a regulatory element in the immune system^4^. However, it’s important to note that IL-4 has a dual nature. On the flip side, it can contribute to fibrosis and exacerbate the pathogenesis of various allergic diseases^6^. While CD4^+^ T cells have long been recognized as a primary source of IL-4, particularly during helminth infections, innate immune cells like basophils and mast cells have been described as significant producers of IL-4^7–9^.

*Litomosoides sigmodontis* is a rodent filarial nematode which serves as a model for human filarial infections, recapitulating immune responses as they occur e.g. in onchocerciasis and lymphatic filariasis patients^10,11^. This filarial nematode gets transmitted via the tropical rat mite *Ornithonyssus bacoti* and migrates through skin and lymphatics to the pleural cavity by 5-8 days post infection (dpi)^12^. In susceptible BALB/c mice these parasites develop to sexual maturity by 35dpi and produce their offspring, the microfilariae, which can be found in the peripheral blood 57dpi onwards. Similar to other helminth infections, *L. sigmodontis* induces a type 2 immune response^13,14^. However, the immunomodulatory capacity of the parasite leads to a strong regulatory immune milieu, facilitating the parasites’ long-term survival in the host. Studies using IL-4 and IL-4R-KO mice have demonstrated a critical role for IL-4 and signaling via the IL-4 receptor in microfilariae control but not adult worm burden^15,16^ Fertility and length of female adult worms were found to be enhanced in the absence of both IL-4 and IL-4R. Additionally, both studies found reduced eosinophil numbers in IL-4 and IL-4R-KO mice and an absence of IgE in infected IL-4 KO mice^15^. In contrast to the susceptible BALB/c strain, C57BL/6 mice eliminate *L. sigmodontis* infection before patency^14,17^. However, in the absence of IL-4, C57BL/6 mice become as susceptible as BALB/c mice^18^. Furthermore, studies have demonstrated the requirement for IL-4 to induce an alternative macrophage phenotype^19–21^, which in turn orchestrates eosinophil-mediated immunity to filarial nematodes^22^.

While many functions for IL-4 have been shown for *L. sigmodontis* infection^11,23^, the contribution of different cellular sources of IL-4 to the infection process has remained unclear. To address this knowledge gap, we employed an IL-4 secretion assay to investigate both innate and adaptive sources of IL-4 throughout the course of *L. sigmodontis* infection. While our experiments confirmed a central role for CD4^+^ T cell-derived IL-4, eosinophils emerged as a major source of IL-4, particularly during the early phase of infection before worms reach sexual maturity. To investigate the impact of eosinophils and their IL-4 contribution on the type 2 response, we utilized dblGATA mice, which lack eosinophils and therefore exhibit eosinophil-derived IL-4-deficiency. Strikingly, while eosinophil deficiency had no discernible impact on macrophage proliferation and polarization, it significantly curtailed the presence of Th2 cells at site of infection, particularly during the middle phase of infection. These findings shed new light on the intricate dynamics of IL-4 sources and on an underappreciated function of eosinophils during *L. sigmodontis* infection.

## METHODS

### Mice

6-8-week-old male and female BALB/c J wild-type (WT) mice were purchased from Janvier Labs (Saint-Berthevin, France). BALB/c Δ*dblGata1* mice were originally obtained from Jackson Laboratory (Bar Harbor, United States), Eotaxin-KO and IL-5-KO mice were originally obtained from Prof. Dr. Klaus Matthaei (Stem Cell & Gene Targeting Laboratory, Australian National University, Canberra, Australia). All mice were bred at the “Haus für Experimentelle Therapie” of the University Hospital Bonn. Mice were housed in individually ventilated cages with unlimited access to food and water and a 12-h day/night cycle. All experiments were performed according to EU directive 2010/63/EU and approved by the Landesamt für Natur-, Umwelt-und Verbraucherschutz (LANUV, Recklinghausen, Germany) under license AZ 81.02.04.2020.A103.

### Natural Infection

Mice were naturally infected with infective *L. sigmodontis* third stage larvae via exposure to the tropical rat mice *Ornithonyssus bacoti* as described before^13^. In short, mice were placed over night in cages with bedding material containing infected mites. The next day, bedding material with the mites was removed and after another 24h mice were moved back to new cages. To ensure comparable infections of both groups, mice from both groups were exposed simultaneously to the same batch of mites containing the infective larvae.

### Parasite recovery

Mice were sacrificed via an overdose of isofluorane (Abbvie, Wiesbaden, Germany) 10, 15, 35/37, 57/60, 72 and 90 days post infection (dpi). To determine adult worm burden, the pleural cavity of individual mice was flushed with 1 ml of PBS (PAA Laboratories, Pasching, Austria) which then contained adult worms. Remaining adult worms in the pleural cavity and the peritoneum were isolated with a dissection probe.

### Isolation of pleural cavity cells

Pleural cavity cells were obtained following lavage with PBS (PAA Laboratories). The first ml of the lavage was collected, worms removed, and cells separated by centrifugation and the supernatant stored at −20°C for cytokine measurements at a later time point. The cells were combined with the cells of a following lavage with 4 ml of PBS.

### Preparation of *L. sigmodontis* antigen

For the preparation of *L. sigmodontis* adult worm extract (LsAg) freshly isolated adult worms were rinsed in sterile PBS before being mechanically homogenized under sterile conditions. Insoluble material was removed by centrifugation at 300 g for 10 min at 4°C. Protein concentrations of crude extracts were determined using the Advanced Protein Assay (Cytoskeleton, Denver, USA).

### Measurement of cytokines and antibodies by ELISA

Cytokine concentrations were determined within the first ml of pleural cavity lavage by ELISA. IL-4, IL-5 and IL-13 (Invitrogen, Massachusetts, USA) were all measured according to kit protocols. To determine parasite-specific immunoglobulin levels in plasma, plates were coated with 10 μg/ml LsAg overnight. After blocking with PBS/1% BSA, plasma was added in serial dilutions. Following incubation, plates were washed and secondary biotinylated antibodies against IgE, IgG1, or IgG2a/2b (BD Biosciences, New Jersey, USA) were added. After washing and incubation with Streptavidin-HRP, plates were washed, TMB substrate added and the enzymatic reaction stopped with sulfuric acid. Optical density was measured at 450 nm (Spectramax 240pc Molecular Devices, California, USA).

### Flow cytometric analyses of pleural cavity cells

Pleural cavity cells were counted using the Countess Cell Counter (Countess Cell Counter, Thermo Fisher, Massachusetts, USA). Cells were then stained for live/dead (Life Technologies) and subsequently incubated with Fc-block (1:500 CD16/CD32 and 1:50 mouse serum) and were then stained with fluorescence-conjugated antibodies. For staining of intracellular cytokines, cells were stimulated for 4 h at 37°C with cell stimulation cocktail containing protein transport inhibitor (eBioscience), then stained with live/dead. After surface antibody staining, cells were fixed for 1h at RT using IC fixation/permeabilization buffer (Biolegend) and cells were then incubated for 20 min at RT in permeabilization buffer (Biolegend). Intracellular staining was performed for cytokines using antibodies for IL-5, IL-13, as well as for GATA3, RELMα, Arginase-1, and Ki67. Samples were analysed by flow cytometry with LSR Fortessa (BD bioscience), data analysed using FlowJo v10.2 software.

### Eosinophil generation for in vitro studies

Bone marrow from naïve mice were used to generate bone marrow-derived eosinophils as previously described^24^. Bone marrow cells were counted using CASY^®^ TT-cell counter system (OMNI life science GmbH & Co. KG) and cells were seeded in in Advanced RPMI medium with 20% FBS, 1% penicillin/streptomycin, 0.1% gentamycin, 2.5% HEPES and 1% Glutamax (Thermo Fisher Scientific GmbH, Germany). Cells were cultured with stem cell factor (SCF) and FMS-like tyrosine kinase 3 ligand (FLT3L) (Peprotech, Rocky Hill, USA) for the first 4 days. Afterwards the growth factors were exchanged with IL-5 (Peprotech, Rocky Hill, USA). Half of the medium was exchanged every other day and on day 8, the cell culture flask was exchanged. After 12 days, cells were harvested and checked for the eosinophil purity using flow cytometry (>95%). Eosinophils were then seeded in 24-well plates and stimulated with LsAg for 24, 48, and 72 h. Supernatant was taken at these time points and stored at -20°C for further use.

### IL-4 secretion assay

To identify IL-4 sources within the pleural cavity, the IL-4 secretion assay kit (Miltenyi Biotech, Bergisch Gladbach, Germany) was employed. Pleural cavity single cell preparations were stimulated for 3 h at 37°C with cell stimulation cocktail containing protein transport inhibitor (eBioscience). Subsequently, an IL-4-specific Catch Reagent was attached to the cell surface of all immune cells. The cells were then incubated for 45 min at 37°C to allow cytokine secretion. The secreted IL-4 bound to the IL-4 Catch Reagent on the secreting cells. These cells were subsequently labeled with a second IL-4-specific antibody, the phycoerythrin (PE)-conjugated IL-4 detection antibody for sensitive detection by flow cytometry for 10 min on ice. Further surface markers for different immune cell populations were added parallel to the IL-4 detection antibody. Cells were then washed and analysed on an LSR Fortessa flow cytometer.

### Statistics

Prism 7.0 (version 7.0c, GraphPad Software) was used for statistical analysis. Differences between experimental groups were assessed by Kruskal-Wallis test for nonparametric data, followed by Dunn’s multiple comparisons test. Data are shown as median with interquartile range.

## RESULTS

### Eosinophils are a major source of IL-4 during *L. sigmodontis* infection

To quantify IL-4 protein levels at the site of infection, pleural lavage was obtained from naïve and infected mice at various time points during the course of *L. sigmodontis* infection, followed by an IL-4 ELISA. IL-4 increased in response to infection and reached a statistically significant elevation by 37dpi (Fig. 1A). Interestingly, during the later stages of infection (57dpi, 72dpi, and 90dpi), time points when microfilariae are present, IL-4 concentrations declined, eventually returning back to levels observed in naïve mice. However, this decline was not due to the clearance of adult worms, as mice still possessed viable worms at 90dpi (Fig. 1B).

**Figure 1:**
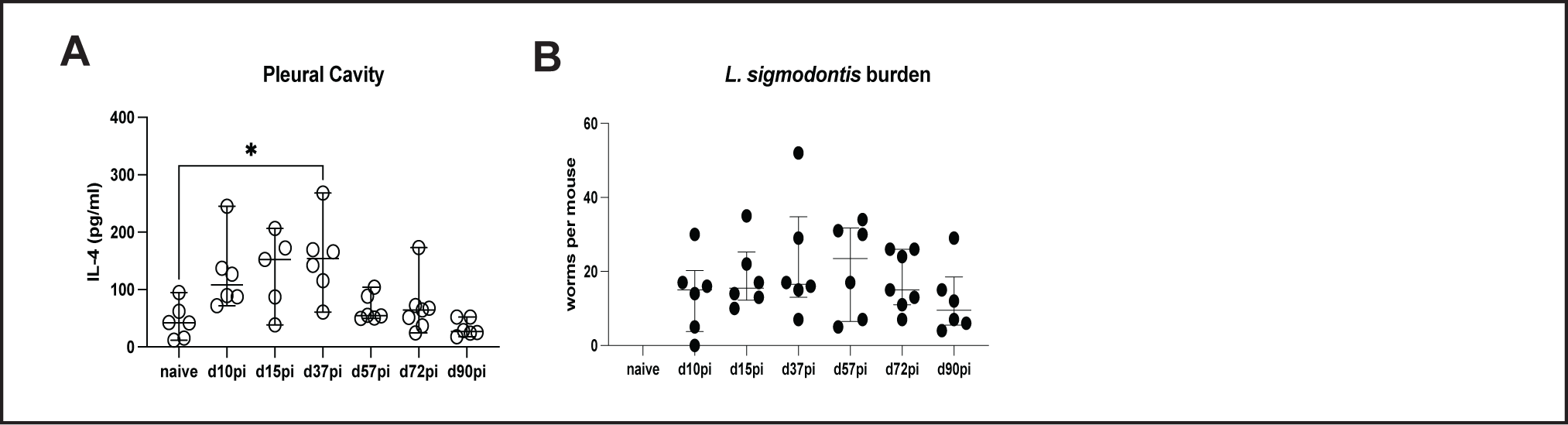
IL-4 is a signature cytokine of *L. sigmodontis* infection. IL-4 protein levels in the pleural lavage of naïve and *L. sigmodontis*-infected BALB/c mice (A). *L. sigmodontis* worm burden during the course of infection (B). Data is representative for 3 independent experiments. Data is shown as median with interquartile range and tested for significances using Kruskal-Wallis-test and Dunn’s post-test comparing each time point to the naïve control (A). *p<0.05

To decipher the sources of IL-4, a murine IL-4 secretion assay kit was employed. In this assay, pleural cavity cells were restimulated and subsequently incubated with an IL-4-specific catch reagent and detection antibody. Cells secreting IL-4 were detected using regular flow cytometric surface staining (Fig. 2A). Similar to IL-4 protein levels, the frequency of IL-4-producing cells significantly increased during the early phase of infection (10, 15 and 37dpi), followed by a decline back to baseline levels during the later stages of infection (Fig. 2B). Similar kinetics were observed for absolute numbers of IL-4-producing cells in the pleural cavity. Notably, the highest influx of immune cells into the pleural cavity occurred at 10dpi, aligning with the peak of IL-4-secreting cell numbers (Fig. 2B). To gain deeper insights into the cellular composition of the IL-4-positive subset among CD45^+^ cells, the immune cells within the IL-4^+^ proportion of CD45^+^ cells were further investigated. As illustrated in Figure 2C, our investigation demonstrates dynamic shifts in the dominant sources of IL-4 at different stages of infection. As expected, CD4^+^ T cells emerged as a consistent and prominent source of IL-4 throughout the course of the infection. However, during the early phases of infection, following the migration of the infective third stage larvae into the pleural cavity and molt into fourth stage larvae, SiglecF^+^CD11b^+^ eosinophils accounted for the highest proportion of IL-4-secreting cells, displaying a decisive role in these initial stages. A surprising discovery was the substantial contribution of Ly6G^+^CD11b^+^ neutrophils to IL-4 production, a role they assumed from their appearance in the pleural cavity at 37dpi, after adult worms developed, persisting until 90dpi, when the natural clearance of the adult worms occurs. In addition to these primary sources, the analysis identified CD11b^+^ cells, which may include basophils and mast cells, as well as CD11b^-^ cells, which may include ILC2, B cells, and NK T cells amongst others (Fig. 2C), cell types which have been described to be involved in *L. sigmodontis* infection. When quantifying the absolute numbers of cells producing IL-4, eosinophils were the most numerous producers of IL-4 during the early time points, followed by CD4^+^ T cells. Again, neutrophils appear during the later stages of infection, when the IL-4 levels in the pleural cavity generally decreased (Fig. 2D). To further confirm that eosinophils are a major IL-4 producer during *L. sigmodontis* infection, ELISA was performed using pleural wash from infected eosinophil-deficient dblGATA mice and wild type (WT) controls. As shown in Figure 2E, dblGATA mice secreted significantly less IL-4 compared to BALB/c WT controls 10dpi (Fig. 2E). Furthermore, bone marrow-derived eosinophils were stimulated with *L. sigmodontis* crude antigen (LsAg) for 24h, 48h, and 72h and IL-4 was measured in the supernatants. IL-4 was detected at all tested time points with the highest production measured after 72h restimulation, while unstimulated eosinophils did not secrete IL-4. This data confirms that eosinophils indeed produce IL-4 in response to filarial antigen in a time-dependent manner (Fig. 2F).

**Figure 2:**
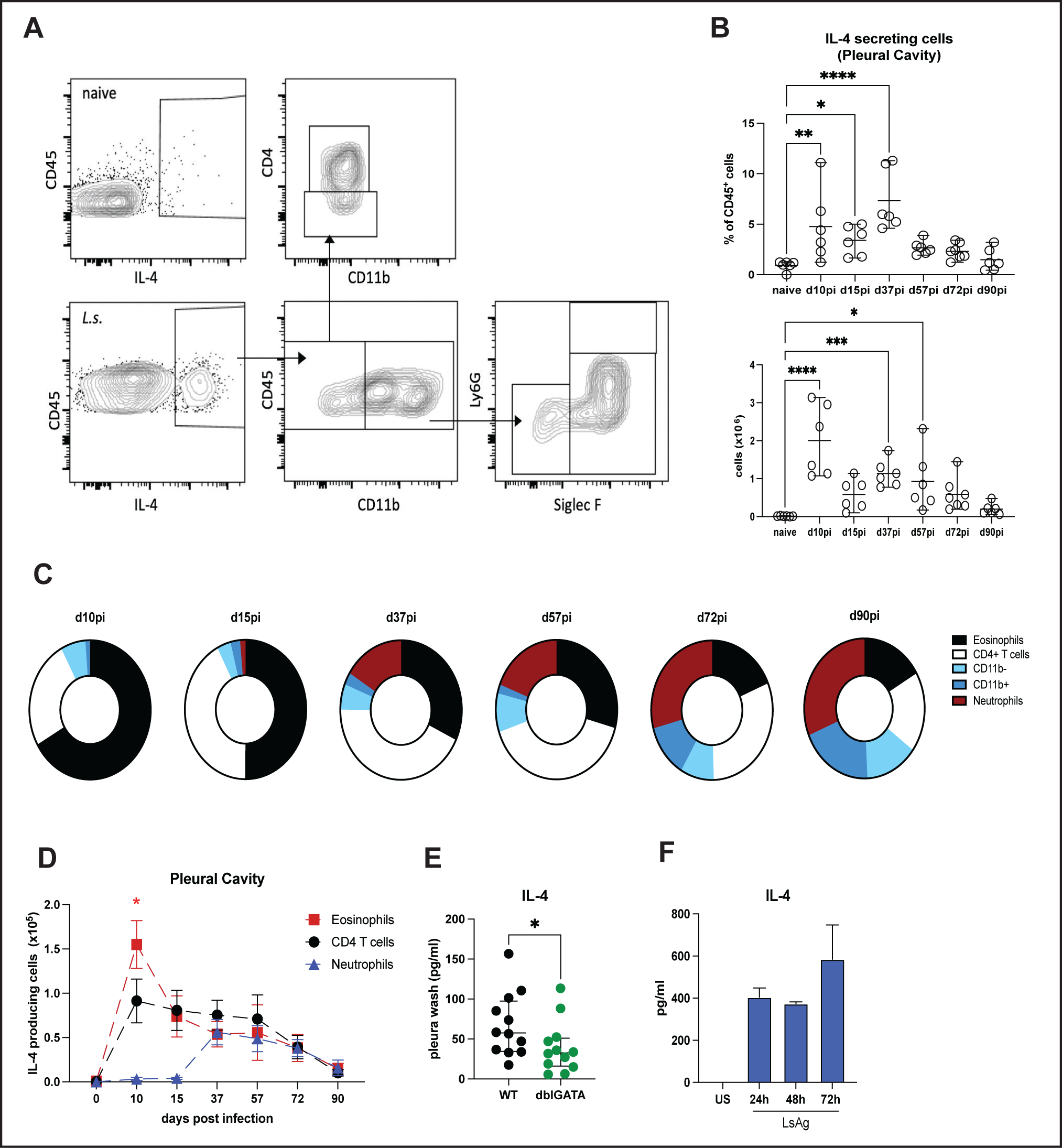
Eosinophils are a major source of IL-4 during filarial infection. Representative FACS-plots displaying gaiting strategy for pleural cavity using the IL-4 secretion assay. Cells are derived from single, live cells. (A). Percentage and absolute number of IL-4^+^ immune cells within the pleural cavity of naïve and *L. sigmodontis*-infected BALB/c mice during course of infection (B). Pie charts representing frequencies of IL-4^+^ cells within the pleural cavity during course of *L. sigmodontis* infection. Each circle represents 100% (C). Absolute numbers of IL-4^+^ eosinophils, IL-4^+^CD4^+^ T cells, and IL-4^+^ neutrophils over the course of infection (D). IL-4 protein levels in the pleural lavage 10dpi with *L. sigmodontis* in BALB/c (WT) and dblGATA mice. IL-4 protein levels measured from supernatants of bone marrow-derived eosinophils stimulated with LsAg for 24h, 48h, and 72h as well as unstimulated controls (F). Data representative for two independent experiments (A-D) or pooled from two independent experiments with n=6 (E, F). Data is shown as median with interquartile range (B, D, E) or as mean ± SEM (F) and tested for significances using Kruskal-Wallis-test and Dunn’s post-test (B, F), two-way-ANOVA with Tukey’s post-test (D) or student’s t-test (E). *p<_0.05, **p_<_0.01, ***p_<_0.001, ****p_<_0.0001.

### IL-4^+^ eosinophils are highly activated and express CCR3, CD125, and ST2 but do not express CD101

Single-cell methodology and transcriptomic studies have largely contributed to uncover new cell subtypes and cell heterogeneity. As such, eosinophils have been characterized in more detail in health and disease in recent years^25–27^. In this study, we employed a flow cytometry approach to elucidate the characteristics of eosinophils on 10dpi, which presented peak eosinophil IL-4 production. At this specific time point of infection, eosinophil numbers are significantly elevated in the bloodstream (Fig. 3A) and at site of infection, the pleural cavity (Fig. 3B). Notably, blood and pleural eosinophilia are dependent on the presence of IL-5 and, to some extent, reliant on eotaxin-mediated recruitment (Suppl. Fig. 1A, B). However, eosinophil numbers do not exhibit a direct correlation with the worm burden at this time point (Fig. 3C). Subsequently, we conducted a comparative analysis of eosinophils in the blood and the pleural cavity of infected mice, focusing on established markers of eosinophils and their activation status. Eosinophils are primarily produced in the bone marrow and are released into the bloodstream upon infection, subsequently infiltrating the pleural cavity, where the parasites reside. Upon entering the pleural cavity, eosinophils upregulate different proteins associated with an either regulatory or inflammatory phenotype as well as increased cell activation compared to their blood counterparts. CCR3 is the receptor for the eotaxins 1 and 2 and is upregulated in pleural eosinophils when compared to blood eosinophils. Similarly, the receptors for IL-33 (ST2) and IL-5 (CD125) were significantly upregulated on eosinophils upon entry into the pleural cavity (Fig. 3D), indicating a requirement for these two cytokines for full activation and induction of effector mechanisms. Recently, CD101^+^ eosinophils were described during infection with the lung-migrating nematode *Nippostronglyus brasiliensis*, exhibiting a more inflammatory phenotype compared to resident CD101^lo^ eosinophils^26,28^. In our setting, a small portion of eosinophils were found to upregulate CD101 when entering the pleural cavity (4,99%; Fig. 3D). Moreover, classical activation markers, including CD69, CD86, and MHCII, exhibited a substantial upregulation specifically in pleural eosinophils when compared to their counterparts in the bloodstream within the same mouse. CD62L was found to be downregulated on pleural eosinophils compared to blood eosinophils. Additionally, these aforementioned markers were assessed and compared between pleural eosinophils from both naïve and *L. sigmodontis*-infected mice at 10dpi and were found to be significantly upregulated in response to infection (Suppl. 1C). Recently, Gurtner et al. presented a protocol to distinguish eosinophil subsets in different tissues using CD80 and PD-L1^25^. However, in our model, eosinophils within the pleural cavity during *L. sigmodontis* infection did not express PD-L1 and exhibited low CD80 expression (data not shown). We subsequently investigated which markers exhibited upregulation specifically on IL-4-producing eosinophils. As shown in figure 3E and F, eosinophils responsible for IL-4 production prominently expressed CCR3 and CD125, with 59.6% of eosinophils displaying ST2 expression. It’s noteworthy that IL-4^-^ but not IL-4-producing eosinophils express CD101 (Fig. 3E & F). CD101 was described to be a feature of an inflammatory phenotype of eosinophils. Additionally, IL-4^+^ eosinophils exhibit a high level of activation, exemplified by their markedly increased expression of CD69 (75%), CD86 (93.8%), and MHCII (53.6%; Fig. 3F).

**Figure 3:**
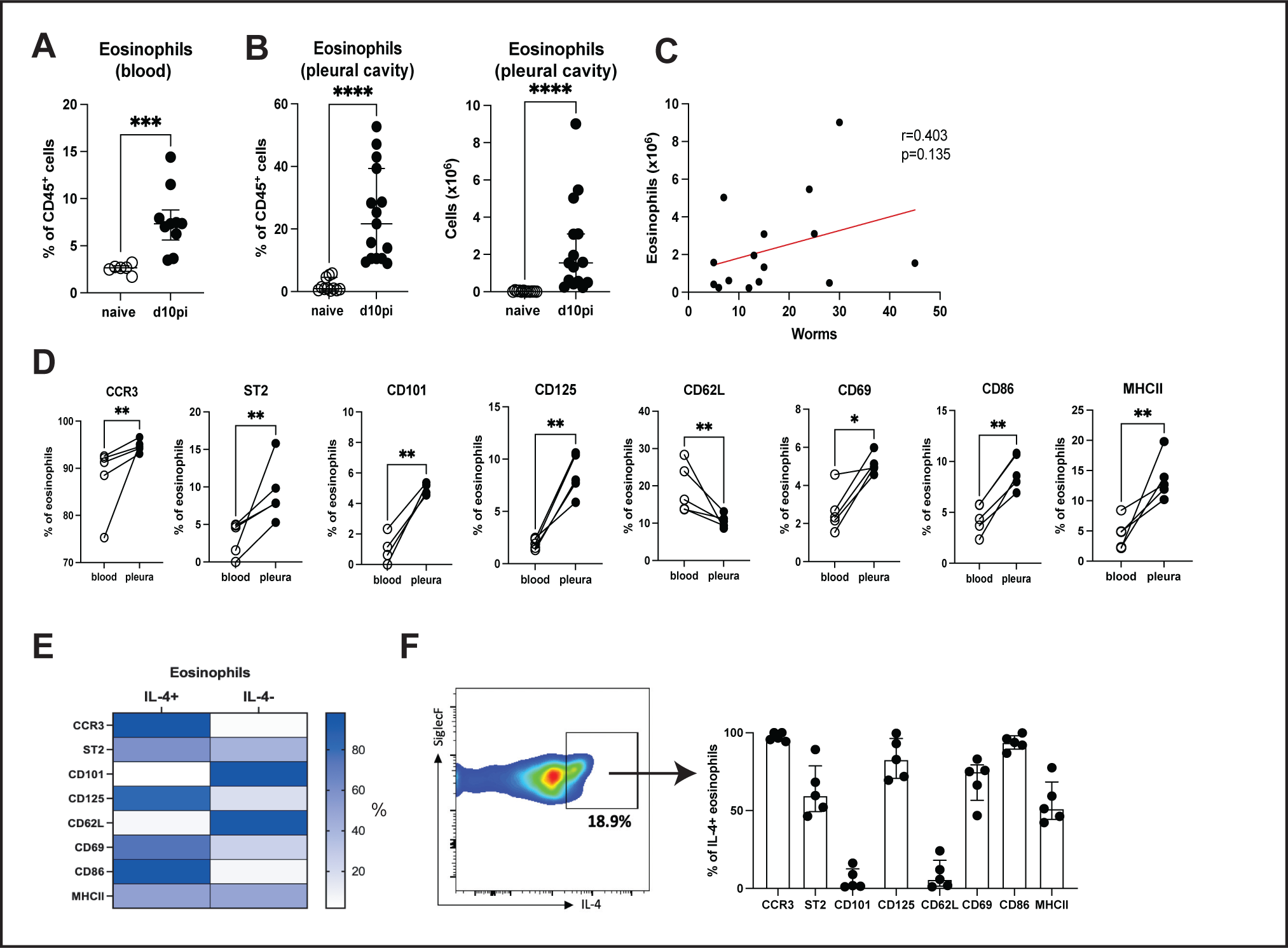
IL-4-producing eosinophils are highly activated and do not express CD101. Eosinophil frequency in blood (A) and pleural cavity (B) as well as absolute eosinophil numbers of pleural cavity in naïve and infected mice 10dpi. Linear regression of worm burden vs eosinophil percentage for 10dpi (C). Comparison of frequencies of CCR3, ST2, CD101, CD125, CD62L, CD69, CD86, and MHCII positive eosinophils in blood and pleura 10dpi (D). Heat map showing the percentage of IL-4^+^ and IL-4^-^ eosinophils expressing indicated activation markers, n=5 per group and marker (E). Representative FACS plot displaying gating for IL-4^+^ eosinophils and graph showing percentage of expression of different markers on IL-4^+^ eosinophils (F). Data shown in A-C is pooled from three independent experiments with 5 mice each. Data shown in D-F is representative for two independent experiments. Data is shown as median with interquartile range (A, B, F). Data was tested for significance using student’s t-test (A, B, D) or tested for correlation using spearman test (C). *p<_0.05, **p_<_0.01, ***p_<_0.001, ****p_<_0.0001

### Eosinophil-derived IL-4 is dispensable for macrophage proliferation and polarization

Having established the significant contribution of eosinophils as a pivotal source of IL-4 during filarial infection, further immune parameters were assessed to understand the impact eosinophils have on the type 2 response. For this, eosinophil-deficient dblGATA mice were used (Fig. 4A). These mice have a complete ablation of the eosinophil lineage, even under conditions, which normally stimulate eosinophil development. As shown in figure 4B, these mice exhibit a significantly increased worm burden during the course of *L. sigmodontis* infection (Fig. 4B). Pleural macrophages are a key cell type during *L. sigmodontis* infection and impairments within the macrophage compartment can have a major effect on infection outcome^23^. The polarization of macrophages to an alternatively activated phenotype is IL-4R-dependent and required for successful elimination of the parasites^23,29^. Comparing dblGATA mice and BALB/c WT controls, no differences were found in frequency and absolute numbers of F4/80^hi^ macrophages in the pleural cavity 15dpi (Fig. 4C & D).

**Figure 4:**
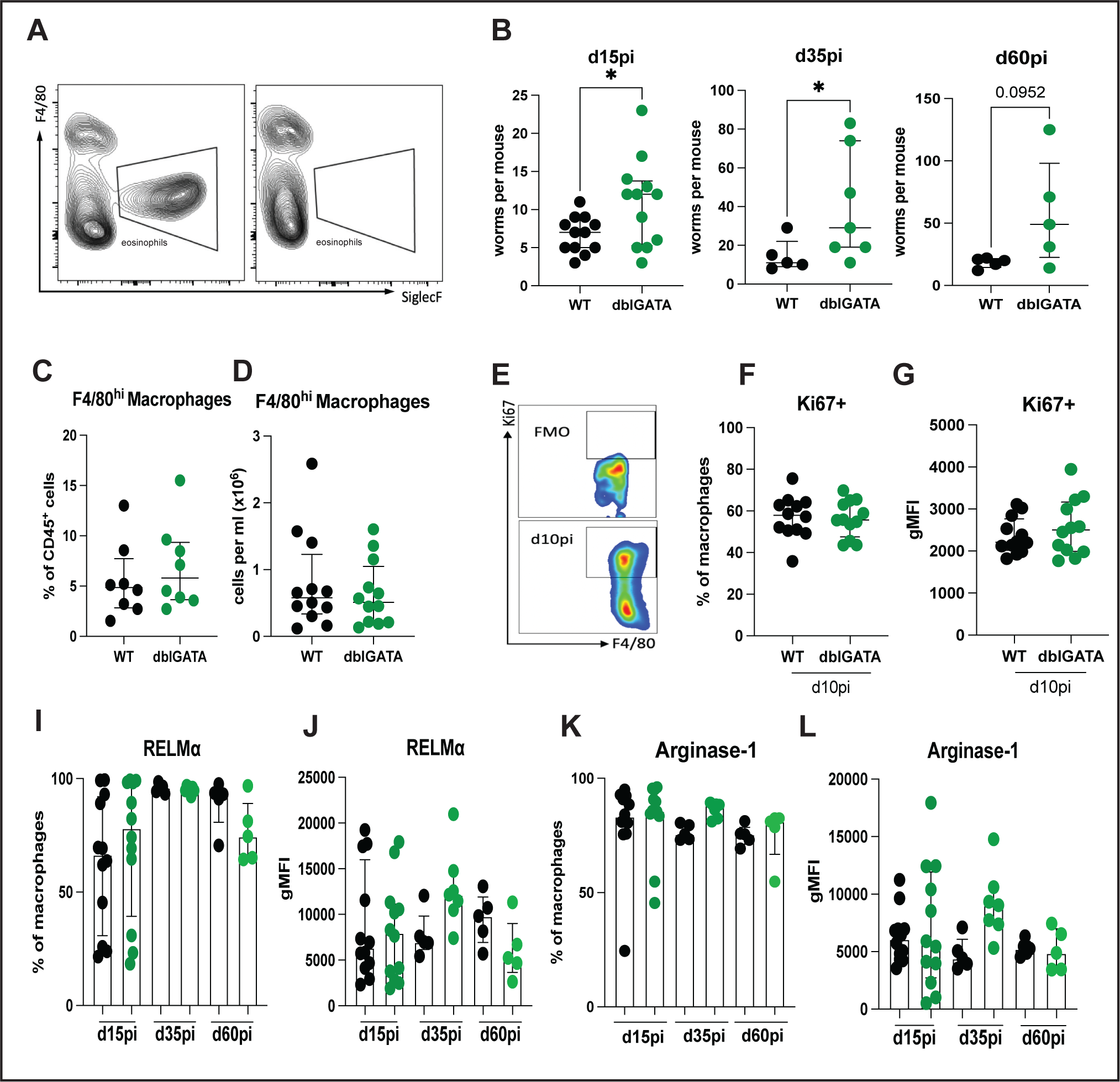
Lack of eosinophils does not affect macrophage polarization and proliferation. Representative FACS plots comparing WT and dblGATA mice for eosinophils, gated as SiglecF^+^F4/80^-^Ly6G^-^CD11b^+^ (A). Worm burden 15dpi, 35dpi, and 60dpi between dblGATA mice and BALB/c WT controls (B). Frequency (C) and absolute number of F4/80^hi^ macrophages in the pleural cavity at 10dpi of dblGATA mice and BALB/c WT controls (D). Representative FACS plot displaying Ki67 staining 10dpi and FMO control for pleural cavity macrophages (E). Percentage (F) and gMFI (G) for intracellular Ki67 staining of pleural F4/80^hi^ macrophages of dblGATA mice and BALB/c WT controls 10dpi. Percentage (I) and gMFI (J) of intracellular RELMα on F4/80^hi^ macrophages between dblGATA mice (green) and BALB/c WT controls (black) 15dpi, 35dpi and 60dpi. Percentage (K) and gMFI (L) of intracellular Arginase-1 on F4/80^hi^ macrophages between dblGATA mice (green) and BALB/c WT controls (black) 15dpi, 35dpi and 60dpi. Data B is pooled from two independent experiments for 15dpi, 35dpi and 60dpi. Data in C, D, F-L is representative for two independent experiments. Data is shown as median with interquartile range (B-D, F, G). Data was tested for significance using student’s t-test (B-D, F, G) or Kruskal-Wallis with Dunn’s post-test (I-L). *p<_0.05.

Macrophages undergo a proliferative burst in the pleural cavity on 10dpi, which depends on the presence of IL-4^29^. Therefore, the expression of the proliferation marker Ki67 was measured on pleural F4/80^hi^ macrophages on 10dpi (Fig. 4E). Eosinophil-deficient mice did not show impaired macrophage proliferation, as frequency of Ki67^+^ macrophages as well as mean fluorescence intensity of Ki67 did not differ between dblGATA mice and WT controls (Fig. 4F & G). Intracellular staining for proteins RELMα (Fig. 4 I& J) and arginase-1 (Fig. 4K & L), both associated with the alternatively activated, IL-4R-dependent phenotype of macrophages did not reveal significant differences during the absence of eosinophils in both frequencies and MFI. A smaller proportion of RELMα+ macrophages was observed in dblGATA mice on d60pi, however this did not reach statistical significance (Fig. 4I). Similar tendencies for reduced MFI for RELMα in dblGATA mice was observed as well (Fig. 4J). Taken together, this data demonstrates that eosinophils are dispensable for macrophage polarization and proliferation in the pleural cavity of *L. sigmodontis*-infected mice.

### Eosinophils are required for an optimal development of the Th2 response

CD4^+^ T cells have been previously described to be essential in controlling worm and microfilariae burden^30,31^. Additionally, depletion of CD4^+^ T cells resulted in diminished Th2 cytokines, eosinophilia and IgE levels, demonstrating the central role for CD4^+^ T cells during filarial infection^30^. The polarization of CD4^+^ T cells to T helper 2 cells requires IL-4. Therefore, differences in CD4^+^ T cells numbers, GATA3 expression - the master transcription factor of Th2 cells - as well as type 2 cytokine production of these T cells were investigated in dblGATA mice and WT controls. CD4^+^ T cell frequency did not differ between both tested groups (Fig. 5A). However, on d35pi absolute CD4^+^ T cell numbers were significantly reduced in dblGATA mice compared to WT controls, indicating an eosinophil-dependent impact on T cell numbers (Fig. 5B). GATA3 expression in percentage and MFI were not changed throughout the infection due to eosinophil-deficiency (Fig. 5 C&D), but absolute numbers of GATA3^+^ CD4^+^ T cells were lower on d35pi (Fig. 5E). Next, we assessed the ability of the GATA3^+^CD4^+^ T cells to produce signature Th2 cytokines. For this, GATA3^+^CD4^+^ T cells were intracellularly stained for IL-5 and IL-13. Frequencies of these cytokines and absolute numbers of IL5^+^ and IL-13^+^ CD4^+^ T cells did not differ between dblGATA mice and BALB/c WT controls 15 and 60dpi, however, 35dpi, CD4^+^ T cells in dblGATA mice produced significantly less IL-5 and IL-13 and absolute numbers of both IL-5^+^ CD4^+^ T cells and IL-13^+^ CD4^+^ T cells being significantly reduced in the absence of eosinophils (Fig. 5 E&F).

**Figure 5:**
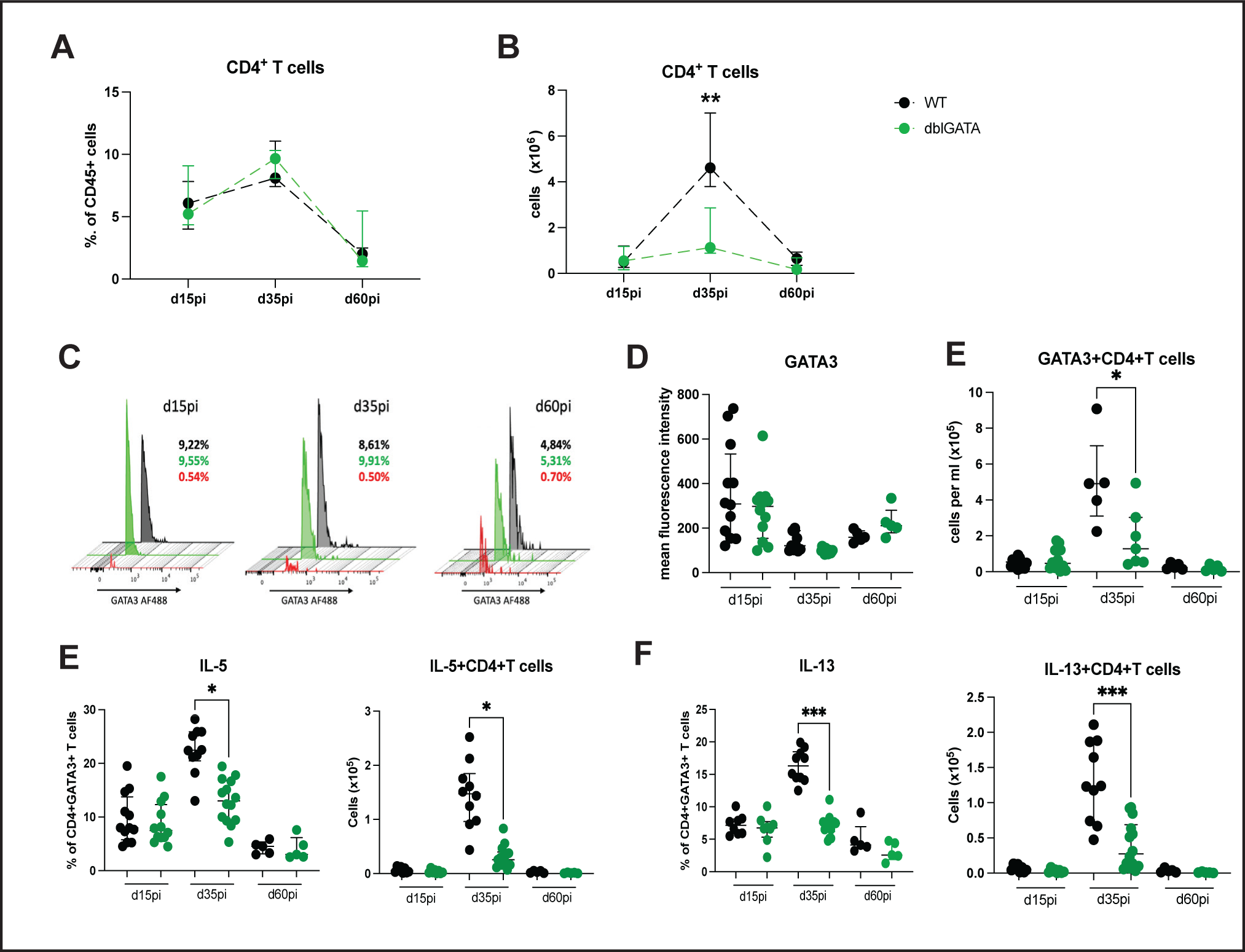
Eosinophils are required for an optimal Th2 response. Frequency (A) and absolute numbers (B) of CD4^+^ T cells in the pleural cavity of dblGATA (green) and BALB/c WT mice (black) 15dpi, 35dpi, and 60dpi. Histograms displaying GATA3 expression between dblGATA (green), WT (black) and FMO controls (red) on pleural CD4^+^ T cells 15dpi, 35dpi, and 60dpi (C). GATA3 gMFI on CD4^+^ T cells (D) and absolute numbers of GATA3^+^CD4^+^ T cells in the pleural cavity of dblGATA (green) and WT controls (black) on 15dpi, 35dpi, 60dpi. Frequency and absolute numbers of intracellular IL-5 (E) and IL-13 (F) on pleural CD4^+^ T cells 15dpi, 35dpi, 60dpi in dblGATA and BALB/c WT controls. Data pooled from two independent experiments, shown as median with interquartile range (A, B, D-F). Data was tested for significance using Kruskal-Wallis with Dunn’s post-test. *p<_0.05, **p_<_0.01.

### Absence of eosinophils has a minimal impact on *L. sigmodontis*-specific antibody production

In line with its function in orchestrating the type 2 response, IL-4 is recognized for its ability for co-stimulating B cells and favoring IgE class switching. However, it remains unclear whether eosinophils and eosinophil-derived IL-4 have an impact on antibody production during filarial infections. To address this question, parasite-specific IgE, IgG2a/b and IgG1 in the serum of infected dblGATA mice and WT controls were measured. Parasite-specific IgE, IgG2a/b and IgG1 were barely detectable 15dpi (Fig. 6 A-C) and no differences were observed between both tested groups. Similarly, no differences in antibody titers for tested IGs were found 35dpi (Fig 6 D-F). On 60dpi optical density (OD) for IgE at 1:2 dilution was significantly reduced in dblGATA mice compared to WT controls (Fig. 6G). However, with lower dilutions, no differences were detected. ODs IgG2a/b was increased in dblGATA mice, but did not reach statistical significance (Fig. 6I). No differences were found for IgG1 (Fig. 6J).

**Figure 6:**
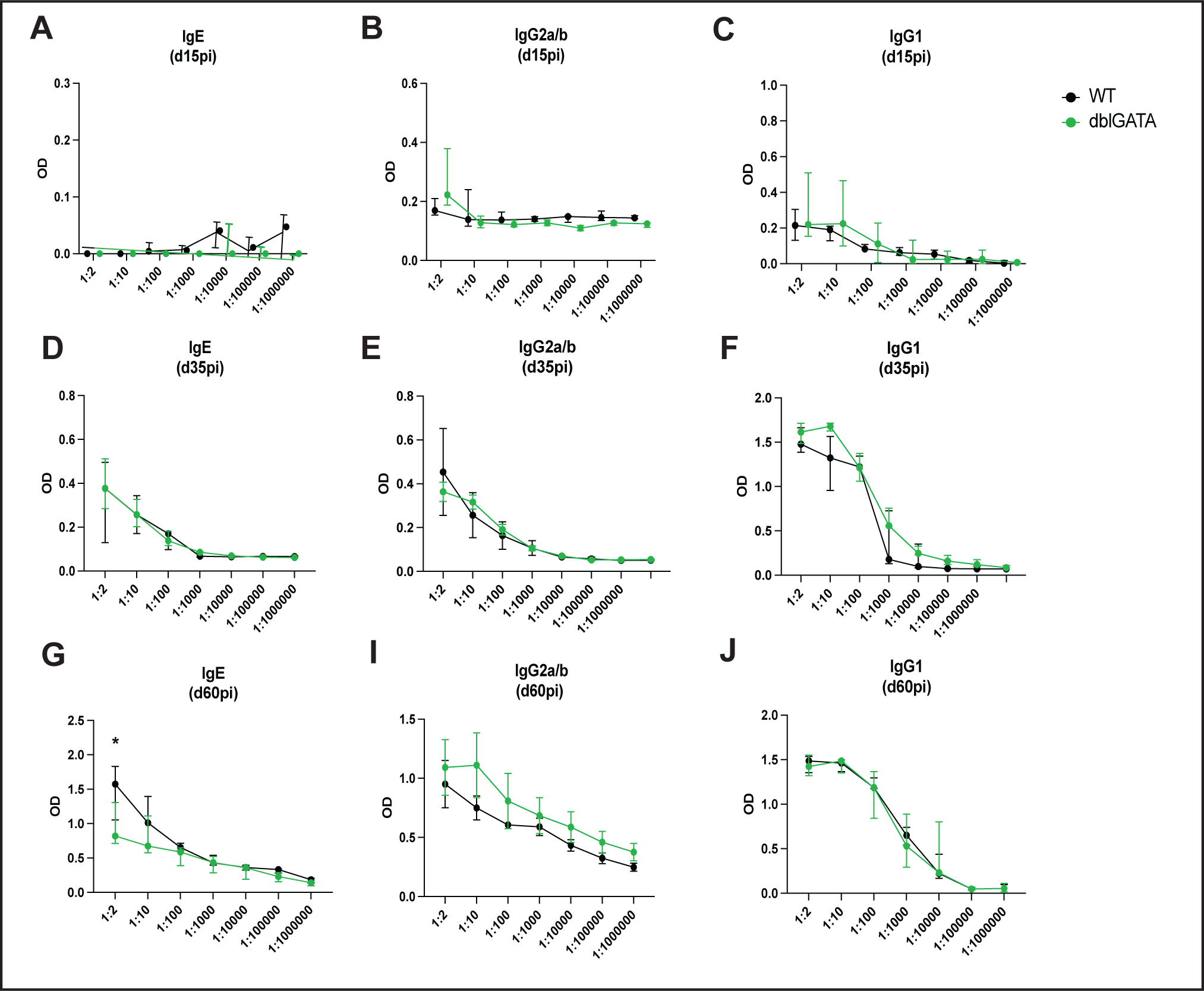
Eosinophil deficiency does not affect antibody production. Optical density (OD) of LsAg-specific IgE (A, D, G), IgG2a/b (B, E, I) and IgG1 (C, F, J) in serum of dblGATA mice and BALB/c WT control 15dpi, 35dpi, 60dpi. Data is pooled from two independent experiments with n=10 per time point. Data is shown as median with interquartile range and were tested for significance with 2-way ANOVA and Sidak’s multiple comparisons test. *p<_0.05

## DISCUSSION

Eosinophils are a hallmark immune cell type of type 2 immunity and highly increased during helminth infections. Eosinophils have been described to have both, protective functions as well as pathology inducing features in filariasis^32^. Specifically, their ability to release granule proteins such as MBP or EPO^33^ as well as the formation of extracellular traps have been investigated in the context of filarial infections^24^. Here, we show that eosinophils are an important source of IL-4, a cytokine central to type 2 immunity. Eosinophils are recruited in high numbers during *L. sigmodontis* infection in the pleural cavity^23^. Our data indicates that these cells heavily contribute to the IL-4 environment within the site of infection. Eosinophils have previously been recognized as a significant source of IL-4 during various disease settings. They have been shown to be major IL-4-expressing cells in white adipose tissue, playing a pivotal role in inducing alternative activation of macrophages^34^. Eosinophils have also been linked to IL-4 production in contexts related to tissue repair, as exemplified in a study on liver regeneration^35^. In a model of a toll-like receptor 2-mediated paw inflammation eosinophils were necessary for “M2-macrophage” polarization and eosinophil depletion resulted in reduced IL-4 levels and increased edema formation^36^. During pulmonary cryptococcosis, eosinophils contribute to IL-4 production and shape the Th2 cell cytokine profile^37^. In human allergic patients, eosinophils may be an important IL-4 source and enhance the allergic response through Th2-formation and inducing isotype switching to IgE^38^. Our data is in line with these studies and show for the first time that eosinophils secrete IL-4 during a filarial infection of the pleural cavity. The eosinophil-derived IL-4 has an impact on the Th2 response and type 2 cytokine production as numbers of CD4+ T cells and IL-5 and IL-13 levels were reduced in the absence of eosinophils. Similar findings were observed in a study of acute schistosomiasis, where lack of eosinophils resulted in lower concentrations of IL-5 and IL-13 in infected liver tissue, indicating that eosinophils participate in the establishment and amplification of Th2 responses^39^. Our data also indicates that the connection between Th2 cells and eosinophils is of major importance to infection outcome and resolution. The eosinophil-induced Th2 cells can in turn not only produce IL-4, but also IL-5, which is necessary for further eosinophil maturation and activation, leading to a positive feedback loop between these two hallmark cell types of type 2 immunity. Additionally, IL-4 and IL-13 production by Th2 cells is necessary for the complex macrophage dynamics during *L. sigmodontis* infection, ultimately leading to worm killing^23^. While an impact of eosinophil deficiency was observed in the CD4^+^ T cell populations, macrophages were not impacted in this study. Our data is hereby in line with two previous studies. In a study by Turner et al, CCR3-KO mice and the *B. malayi* implant model were used to demonstrate that lack of CCR3-dependent eosinophilia did not impact alternative activation of macrophages as well as RELMα and arginase-1 expression^22^. However, lack of eosinophils resulted in an impairment in RELMα secretion. Similarly, Jackson-Jones et al did not observe an eosinophil-dependent impairment in macrophage proliferation using dblGATA mice but saw a reduction in RELMα expression in their model of in vivo IL-33 delivery^40^. However, both these studies have investigated the peritoneal cavity, so tissue-depending differences in RELMα production and secretion could be an explanation for the different study outcomes. The question remains why eosinophil-derived IL-4 does not impact macrophages, but CD4+ T cell-derived IL-4 does. Depleting CD4+ T cells during *L. sigmodontis* infection resulted in failure of macrophages to proliferate in absence of Th2 cells^23,41^. Importantly, IL-13 is capable of inducing macrophage proliferation as well and may be taking over a more dominant, compensatory role during the absence of eosinophil-derived IL-4. Another intriguing observation of this study was the finding that neutrophils are also capable of producing IL-4 during a type 2 immune response. Neutrophils have been described to be involved in helminth infections in general^42^ and in filarial infection in particular^43^. However, their precise role and how they influence various facets of the type 2 immune response remain elusive. Future studies will investigate the role neutrophils have in the context of type 2 responses.

A notable limitation of this study is the reliance on dblGATA mice as opposed to an eosinophil-specific IL-4 KO mouse model. To our knowledge, such a model does not presently exist, rendering dblGATA mice the most suitable option for investigating eosinophils and their absence in this context. Furthermore, while our study primarily focuses on IL-4, we cannot dismiss the possibility that eosinophils might exert their effects on the Th2 response through other means. Previous research has highlighted the T cell-modulating and -regulating capabilities of neutrophil-derived granule proteins^44^, implying a similar potential for eosinophils in shaping the immune response. These aspects warrant further investigation to comprehensively understand the mechanisms at play in eosinophil-mediated immune modulation.

In summary, our study presents for the first-time eosinophils as a major source of IL-4 during filarial infections and this IL-4 is necessary for Th2 cell development and cytokine production during the middle phase of infection.

## Acknowledgements

We would like to acknowledge the assistance of the Flow Cytometry Core Facility at the Institute of Experimental Immunology, Medical Faculty at the University of Bonn.

## Statement of Ethics

All experiments were performed according to EU directive 2010/63/EU and approved by the Landesamt für Natur-, Umwelt-und Verbraucherschutz (LANUV, Recklinghausen, Germany) under license AZ 81.02.04.2020.A103.

## Author contributions

Conceptualization: MPH, JA

Data curation: CG, JA.

Experimentation: CG, PPS, AV, HB, HB, MK, FR, BL, JA.

Investigation: CG, PPS, AV, HB, HB, MK, JA.

Resources: AH, MPH.

Supervision: AH, MPH, JA.

Writing - original draft: CG, MPH, JA.

Writing - review & editing: all authors

## Data availability statement

Data that support the interpretation of findings presented in this paper are openly available. Enquiries can be directed to the corresponding authors.

---

**Supplement figure 1:**
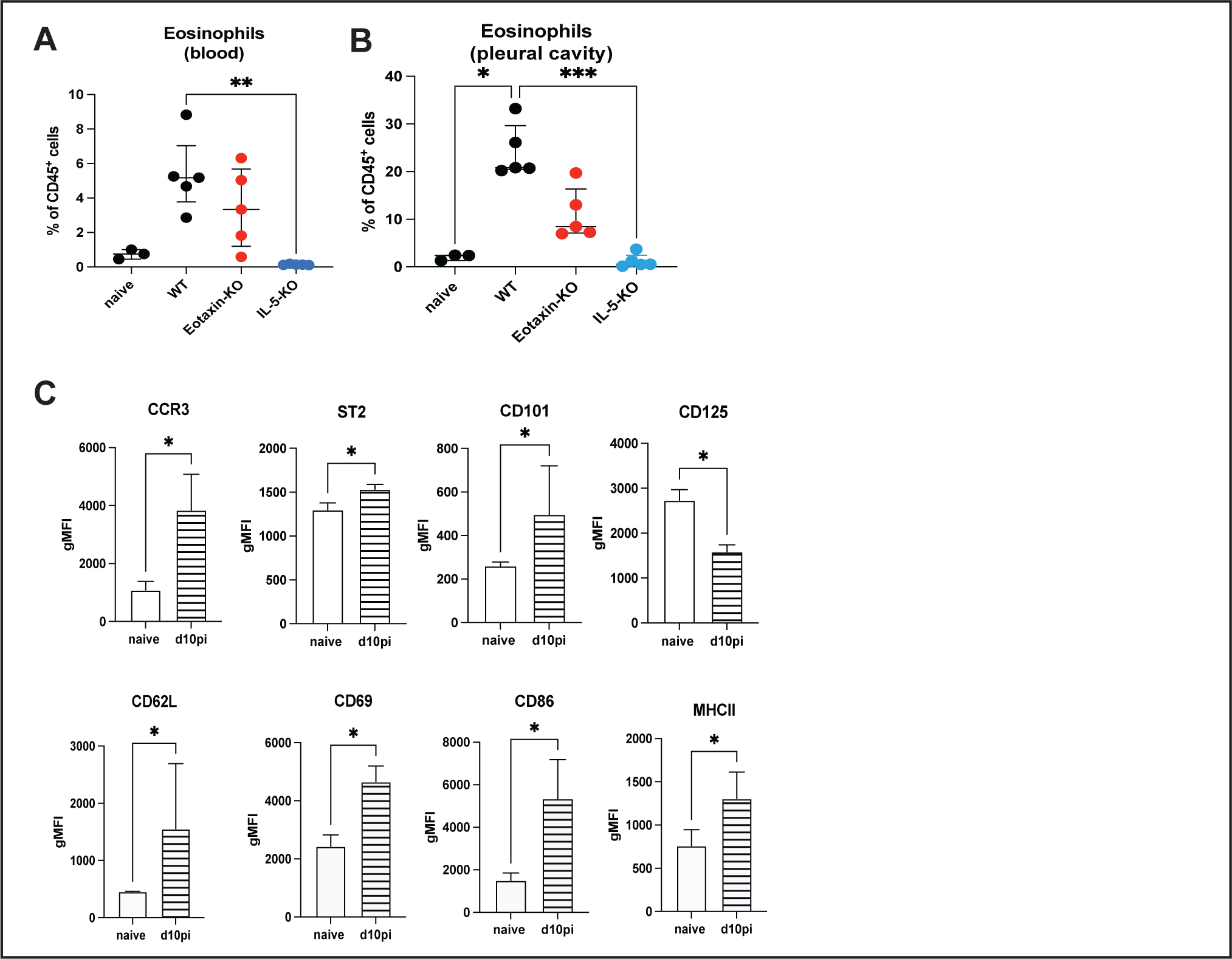
Eosinophil frequencies in blood (A) and pleural cavity (B) of naïve mice, BALB/c controls as well as eotaxin-KO and IL-5-KO mice 10dpi. gMFI of different eosinophil markers and activation markers on pleural eosinophils of naïve and infected BALB/c WT mice d10pi (C). Data from one experiment (A, B) or representative for two independent experiments with n=3 for naive group and n=7 for infected group (C). Data is shown as median with interquartile range and tested for significances using Kruskal-Wallis with Dunn’s post-test. *p<0.05, **p<0.01, ***p<0.001.

